# Evolution in a moving frame of reference

**DOI:** 10.1101/2021.09.20.461092

**Authors:** Nikunj Goel

## Abstract

The study of range expansion has produced a new class of directional and non-directional expansion models of evolution, representing spatial sorting and gene surfing, that are widely considered spatial analogs of selection and drift, respectively. However, unlike classical models of selection and drift that measure evolutionary change within a population across generations, expansion models measure phenotypic change in a moving frame of reference by tracking successive generations of individuals occupying the expansion front. This raises two fundamental questions: what drives evolution in a moving frame of reference, and whether spatial sorting and gene surfing are truly spatial analogs of selection and drift. We answer these questions by using Price’s theorem to analyze evolution in a moving frame of reference. We find that in expansion models, evolutionary change is driven by heritable variation in parental phenotypes and differential sorting fitness—the number of offspring a parent leaves in a newly colonized patch. This finding implies that gene surfing is a spatial analog of drift, but spatial sorting, as currently defined, is not a spatial analog of selection.

## Introduction

Classical evolutionary theory, developed over the 20th century, has profoundly shaped our conceptual understanding of the mechanisms driving evolutionary change in natural systems (Mayr and Provine 1998; Provine 1971; Provine 1977). The theory formalized the verbal arguments of Darwin and Wallace (1858) into mathematical models that can predict the causes and consequences of phenotypic variation. These classical models fall into two broad groups: directional and non-directional models. Directional models invoke some form of selection driven by a causal mechanism linking heritable phenotypic variation to an organism’s lifetime reproductive success (Crow and Kimura 1970; Falconer 1960; Fisher 1930; Maynard Smith 1982). Non-directional models, on the other hand, produce evolutionary change through random variation in reproductive output (Wright 1931).

Over the past three decades, studies of population expansion in field (Duckworth 2008; Forsman et al. 2011; Hanski et al. 2004; Henn et al. 2016; Hill et al. 1999; Lombaert et al. 2014; MacKay and Lamb 1979; Peroni 1994; Phillips et al. 2006; Willi et al. 2018) and in laboratories (Bosshard et al. 2017; Ochocki and Miller 2017; Szűcs et al. 2017; Usui and Angert 2024; Weiss-Lehman et al. 2017; Williams et al. 2016) have led to the development of a new class of evolutionary models that mirror this dichotomy (Miller et al. 2020). The directional models describe spatial sorting, in which phenotypes are filtered at the expansion front based on heritable variation in dispersal-related traits, even when those traits do not confer a fitness advantage (Phillips et al. 2010; Shine et al. 2011; Travis and Dytham 2002). Non-directional models describe gene surfing, in which phenotypes can rise in frequency along the invasion vanguard due to random sampling of founding populations during serial colonization events (Edmonds et al. 2004; Hallatschek and Nelson 2008; Klopfstein et al. 2006). Because of these parallels, some regard spatial sorting and gene surfing as directional and non-directional spatial analogs of selection and genetic drift, respectively (Peischl and Gilbert 2020; Phillips and Perkins 2019; Slatkin and Excoffier 2012).

Despite these apparent similarities, expansion models fundamentally differ from their classical counterparts in how they measure evolutionary change. Classical models of selection and drift quantify evolutionary change in a closed population by observing the difference in mean phenotype between offspring and parents within a static arena (Crow and Kimura 1970; Fisher 1930; Maynard Smith 1982). By a static arena, we mean a non-permeable, bounded geographical region that encompasses parents and all their progeny. Expansion models, in contrast, measure evolutionary change in a moving frame of reference by tracking the difference in mean phenotype between successive generations of individuals occupying the expansion front (for example, see Peischl and Gilbert (2020) and Phillips and Perkins (2019)). These differences raise several questions. What drives evolution in a moving frame of reference? Are these drivers different from those that determine evolution in a static arena? Are models of selection and drift truly analogous to models of spatial sorting and gene surfing? If so, how can we demonstrate these analogies in a general setting rather than through model-by-model comparisons?

Our limited ability to answer these questions partly stems from how current expansion models of evolution are developed (Goel et al. 2025; Peischl and Gilbert 2020; Phillips and Perkins 2019). The modeler starts with a comprehensive set of simplifying assumptions relevant to a particular biological system—such as the nature of the phenotype, the mechanism of genetic inheritance, and the relationships linking phenotype to reproduction and dispersal—and then analyzes these assumptions to derive mathematical results describing evolutionary change. Consequently, this ‘all assumptions first’ approach may obscure the general principles driving evolution because the intuition built for one model may not carry over to other models with different underlying assumptions, making it difficult to compare classical and expansion models of evolution in a general manner.

Recognizing these limitations, some theorists have argued for a different approach for conceptualizing and building a mathematical theory of evolution (chapter 6 in Rice 2004; Rice 2020). Rather than laying out all relevant assumptions at the start, one first defines a minimal set of assumptions that applies to a broad class of biological systems. The mathematical formula derived from this small set of assumptions then serves as the foundational result for quantifying evolution, from which special-case models can be recovered by imposing additional assumptions. Building theory this way can reveal an underlying common thread uniting otherwise disparate models and provide a broad conceptual understanding of the drivers of evolution. In evolutionary theory, this foundational result is Price’s theorem (Price 1970; Price 1972).

At its core, Price’s theorem is an accounting identity that maps the relationship between phenotypes of ancestors in the ancestor set and their descendants in the descendant set over a span of time:

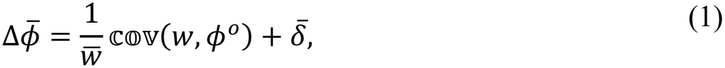

where 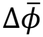 is the change in mean phenotype between descendants and ancestors, *w*_*i*_ is the number of descendants of ancestor *i* in the descendant set, 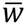 is the mean number of descendants an ancestor leaves in the descendant set, 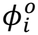 is the mean phenotypic value of descendants of *i* in the descendant set, and 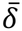 is the remaining phenotypic change that is not covered by the covariance term. The usefulness of Price’s theorem stems from its generality: the theorist is free to choose the identities of ancestors and descendants, as well as the mapping relationship connecting entities in these two sets. Price’s theorem then dictates what must hold irrespective of these choices (Frank 2012).

With only a few additional assumptions, one can use Price’s theorem to gain conceptual insights about the drivers of evolutionary change in a static frame of reference (i.e., selection and drift). Suppose that ancestors and descendants are parents and offspring in a closed population, respectively, with *ϕ*_*i*_ denoting the phenotype of the *i*th parent and 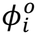 the mean phenotype of its offspring. Under these assumptions, the term *w*_*i*_ in Price’s theorem represents the lifetime reproductive success, *W*_*i*_, of the *i*th parent and yields the following evolutionary change:

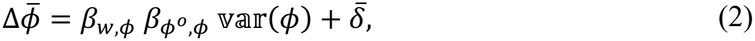

which mathematically encodes the verbal argument of Darwin and Wallace (1858) (Gardner 2008). Here, 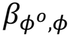— the regression coefficient of offspring phenotype 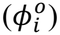 on parental phenotype (*ϕ*_*i*_)— represents heritability; var(*ϕ*) measures variation among parental phenotypes; and *β*_*w,ϕ*_—the regression coefficient of relative fitness 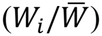 on parental phenotype (*ϕ*_*i*_)—captures differential lifetime reproductive success. The remaining term 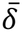 is the population-wide transmission bias, which many evolutionary models assume to be zero. By supplying additional simplifying assumptions, one can derive analytical expressions for regression coefficients and variance terms and recover standard models of selection in classical evolutionary theory (Frank 1997; Luque 2017; Rice 2004).

In a static frame of reference, Price’s theorem also admits an alternative, but equally valid, interpretation. Because Price’s theorem makes no assumption about *why* parental phenotype is associated with relative fitness, it also accommodates genetic drift: in finite populations, the association, that is, *β*_*w,ϕ*_, can arise purely by chance (Okasha 2006; Rice 2004). Price’s theorem thus unifies both the directional and non-directional classical models within a single mathematical identity. For these reasons, Price’s theorem is widely regarded as a general mathematical descriptor of evolution and has thus been crowned by some as the fundamental theorem of evolution (Queller 2017).

However, we do not know if Price’s theorem also describes directional and non-directional evolution in a moving frame of reference. Addressing this gap can help us answer fundamental questions outlined above and unify classical and expansion models of evolution under one umbrella. Our paper has four goals that address this knowledge gap. First, we propose a general mathematical equation of evolution that describes how evolution ensues in a moving frame of reference. We call this equation the *sorting theorem* and show it to be a special case of Price’s theorem. Second, by supplying a few basic assumptions, we use the sorting theorem to identify drivers of directional and non-directional evolutionary change that transcend any particular special-case expansion model of evolution. Third, using the sorting theorem as a springboard, we derive six special-case models of evolution. Five of these have appeared independently in the literature (Goel et al. 2025; Peischl and Gilbert 2020; Phillips and Perkins 2019; Slatkin and Excoffier 2012), but the sorting theorem reveals these expansion models as instances of a common framework, exposing the shared mathematical structure that has so far gone unrecognized. Finally, in the Implications section, we use Price’s theorem to juxtapose drivers of evolution in a static and moving frame of reference to evaluate whether spatial sorting and gene surfing are, in fact, directional and non-directional spatial analogs of selection and genetic drift, respectively.

### Sorting theorem

Consider two groups, labeled as *x* and *x*′ (see Fig. 1). At time *t*, a population of *N* ancestors occupy group *x*, while group *x*′ is empty. We use *ϕ*_*i*_ to denote any numerically quantifiable phenotype of ancestor *i*, and 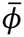 to denote the mean phenotype of the ancestors. At a later time *t′*, a fraction *V*_*i*_ of ancestor *i*’s *W*_*i*_ descendants are in group *x*′. To track the residency of *i*’s descendants at time *t′*, we define an indicator function 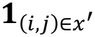, which is one if the *j*th descendant of *i* is in group *x*′, and zero otherwise. We refer to the product *V*_*i*_*W*_*i*_, the number of descendants of *i* in group *x′*, as the *sorting fitness* of *i*, and it is given by the sum of indicator functions over *i*’s descendants:

**Figure 1:**
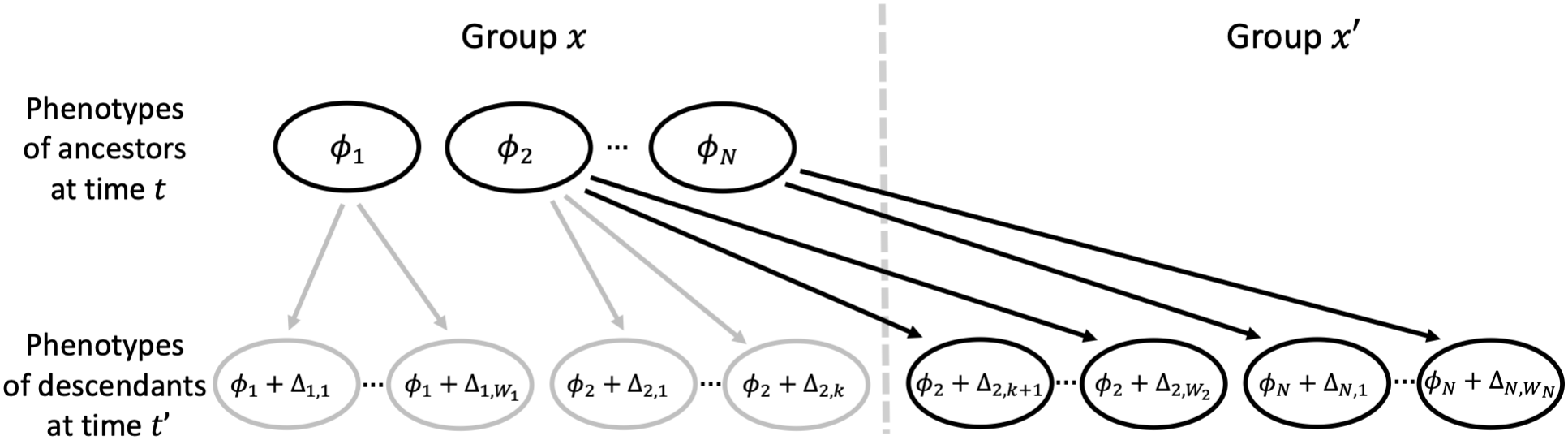
A schematic diagram showing the relationship between ancestors in group *x* at time *t* (the ancestor set) and their descendants in group *x′* at time *t′* (the descendant set). Here, the sorting fitness of the 1st, 2nd, and *N*th ancestors is 0, *W*_2_ − *k*, and *W*_*N*_, respectively.

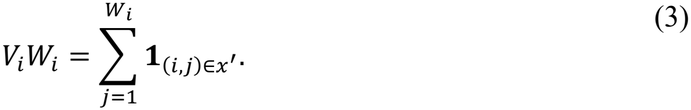

Assuming the phenotype of the *j*th descendant is Δ_*i,j*_ more than its ancestor *i*, the mean phenotype of *i*’s descendants in group *x*′ is

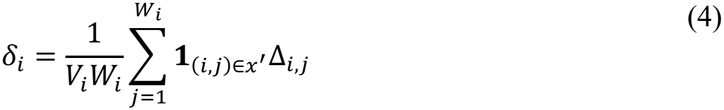

more than *ϕ*_*i*_. Based on this notation, the mean phenotype of the descendants in group *x*′ at time *t′* is

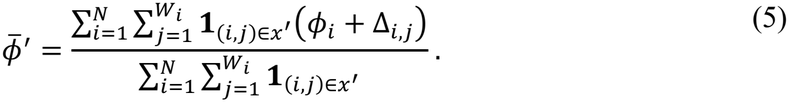

We obtain the above equation by adding the phenotypic values of all descendants in group *x*′ and dividing by the total number of descendants in group *x*′. Using equations (3) and (4), we re-express equation (5) as

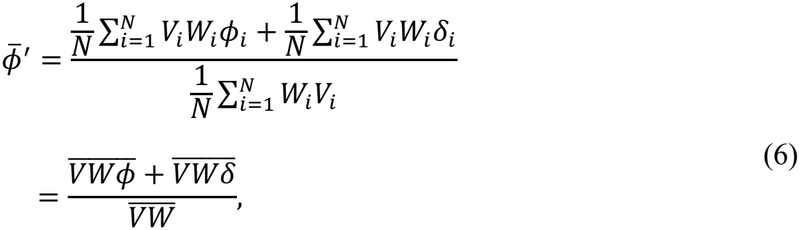

where 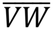, referred to as the *mean population sorting fitness*, is the mean number of descendants an ancestor leaves in group *x′*. Here, we use the bar to denote the mean. Using the covariance identity, 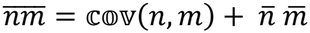, we get

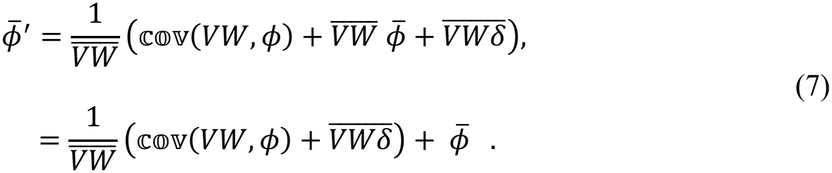

Subtracting 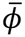 from both sides, we find that the mean phenotypic change between descendants in group *x′* at time 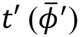 and ancestors in group *x* at time 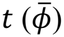 is

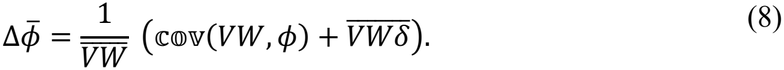

By applying the covariance identity to the second term, equation (8) can be re-expressed as

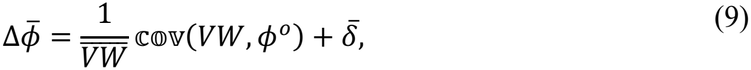

where 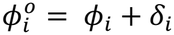 is the mean phenotype of the descendants of *i* in group *x′* at time *t′*. We refer to equation (9) as the *sorting theorem* and consider it as a general mathematical descriptor of evolution in a moving frame of reference (see the special-case models section below).

Sorting theorem (Eq. 9) bears a remarkable resemblance to Price’s theorem (Eq. 1). This is because the sorting theorem is a special case of Price’s theorem. Since a theorist is free to choose the identity of ancestors and descendants in Price’s theorem, let’s define the ancestor set as the ancestors in group *x* at time *t* and the descendant set as the descendants in group *x′* at time *t′* (Fig. 1). Based on these definitions, the term *w*_*i*_—the number of descendants of ancestor *i* in the descendant set—in Price’s theorem (Eq. 1) is equal to the sorting fitness of ancestor *i* (*V*_*i*_*W*_*i*_), yielding the sorting theorem.

#### Mechanisms driving evolution in a moving frame of reference

To build intuition for what drives evolutionary change in a moving frame of reference, we supply some basic assumptions to the sorting theorem that apply to many biological systems of interest. We consider that ancestors are parents and descendants are offspring, such that *t′* − *t* is equal to one generation, and group *x* and *x′* represent occupied and unoccupied spatial units, such as patches in a metapopulation or an invasion front. The sorting theorem tells us that in a moving frame of reference, evolution ensues by any biological process that leads to covariance between the number of offspring a parent leaves in the newly colonized patch *x′* (i.e., sorting fitness *V*_*i*_*W*_*i*_) and the mean phenotype of those offspring. The sorting theorem includes an additional term 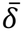, which captures mean transmission bias between parents and their offspring in patch *x′*, which can be assumed to be zero in many biological scenarios (see the special-case models in the section below).

More importantly, the sorting theorem does not specify what biological processes produce the covariance between sorting fitness and offspring phenotypes. One possible explanation for the covariance is that both offspring phenotype and sorting fitness are associated with parents’ phenotypes (Fig. 2). Offspring resemble their parents because of the laws of genetic inheritance, and parents’ phenotypes are causally linked to the individual’s reproductive success and dispersal capacity. Thus, the parents’ phenotype is a common factor that leads to covariance between offspring phenotypes and sorting fitness. To see this more clearly, suppose the parental phenotype *ϕ*_*i*_ determines both sorting fitness (*V*_*i*_*W*_*i*_) and offspring phenotype 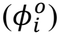. Then we can expand the covariance in the sorting theorem to show that

**Figure 2:**
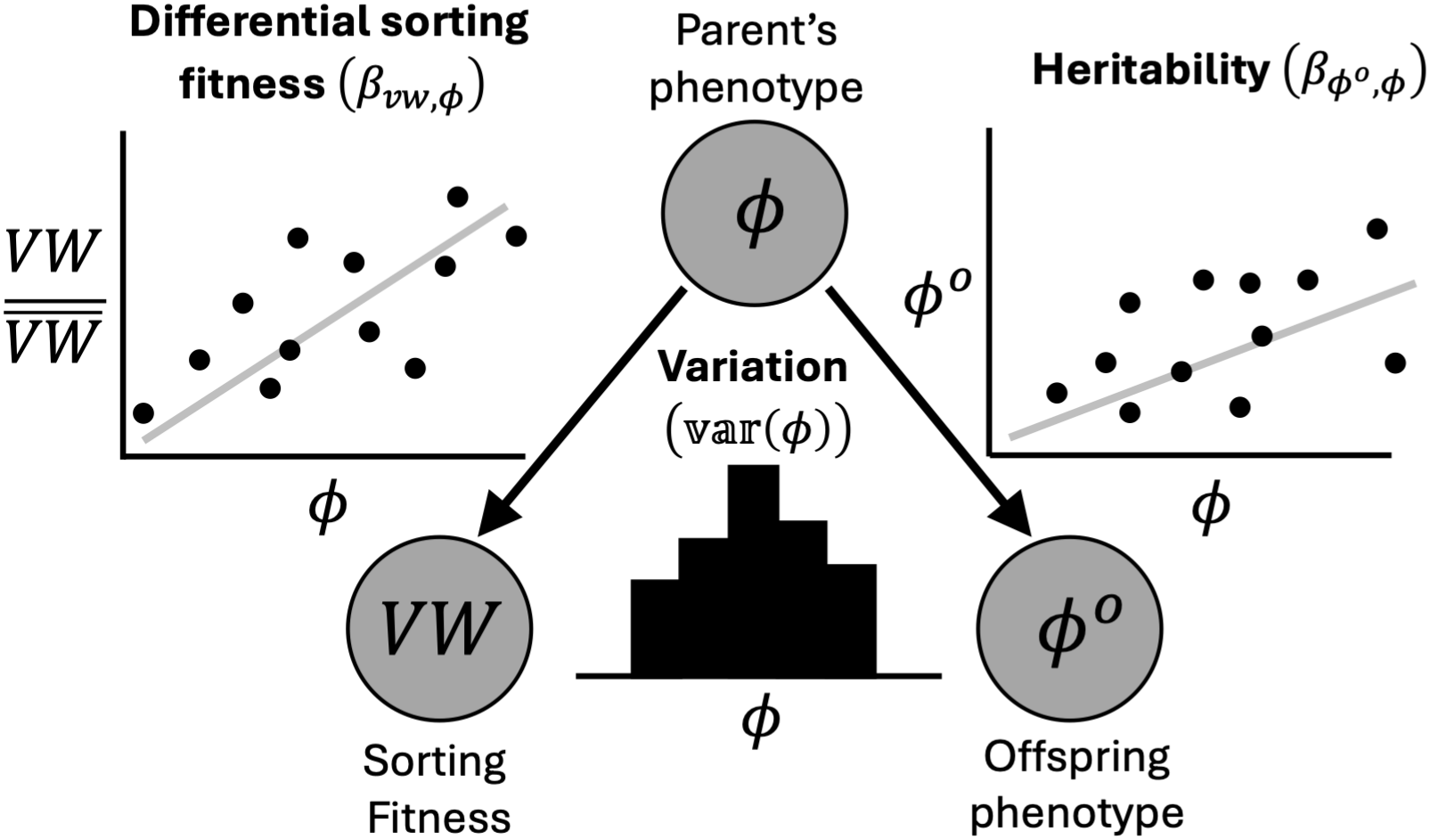
The sorting theorem suggests that in a moving frame of reference, evolution ensues due to statistical association of parents’ phenotypes (*ϕ*_*i*_) with relative sorting fitness 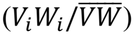 and mean offspring phenotype 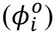. These associations can be explained by laws of genetic inheritance 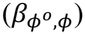 and a causal biological mechanism that links an individual’s phenotype with its reproduction and dispersal (*β*_*vw,ϕ*_). However, the association between parents’ phenotype and sorting fitness could also arise because of random chance.

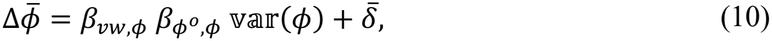

where *β*_*vw,ϕ*_ is the regression coefficient that captures the causal link between parent’s phenotype (*ϕ*_*i*_) and *relative sorting fitness* 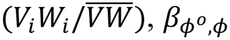 is the regression coefficient that captures the laws of genetic inheritance, and var(*ϕ*) captures variation in phenotypes of parents (Fig. 2). We retain the term 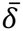 for completeness. This leads us to a simple biological intuition: in a moving frame of reference, directional evolutionary change is driven by heritable variation in phenotypes that confers differential sorting fitness to parents.

However, alternative biological mechanisms can also produce a statistical association between parental phenotype and sorting fitness. In our discussion above, we assumed this association is causal, as the parental phenotype directly influences reproduction and dispersal. The sorting theorem, however, makes no claim that the association (*β*_*vw,ϕ*_) is causal. For example, sorting fitness is a random outcome whose realized value will generally differ from its expectation because reproduction and dispersal are stochastic processes. This stochasticity can produce a non-zero *β*_*vw,ϕ*_ purely by chance, even when all parents share the same expected sorting fitness, resulting in non-directional evolutionary change.

These results illustrate the usefulness of the sorting theorem in building intuition about the drivers of evolution in a moving frame of reference. We make only basic assumptions about the quantities involved (offspring phenotype, parental phenotype, and number of offspring and their location) and how they might be biologically related (the inheritance mechanism, 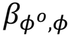, and the link between phenotype and sorting fitness, *β*_*vw,ϕ*_), without specifying the precise mathematical relationships. Yet the sorting theorem dictates what must hold among these quantities, enabling biological inferences that transcend any particular system.

### Special-case models

In this section, we make additional assumptions to derive six special-case models, five of which have been derived independently elsewhere. These models include four population genetic models (two directional and two non-directional), in which we understand how genes relate to phenotypes, and two quantitative genetic models (both directional), in which we treat phenotypes without explicit reference to underlying genes. We carry over the assumptions we used to derive equation (10). For details about the analytical calculations, see the supplementary information.

#### Haploid model of gene evolution

Consider an infinitely large population of haploid parents with genotypes *A* and *a* in frequency *p* and 1 − *p*, respectively. We define *i*’s phenotype as

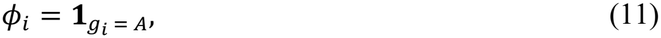

which equals one if the *i*th parent has genotype *g*_*i*_ = *A* and zero if *g*_*i*_ = *a*. By construction, 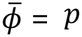, such that the evolutionary change Δ*p* is the difference between the frequency of allele *A* among offspring in patch *x′* and 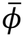. Assuming perfect transmission (no mutation, so 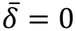 and 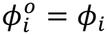), and denoting the sorting fitness of parents with genotypes *A* and *a* by *V*_*A*_*W*_*A*_ and *V*_*a*_*W*_*a*_, respectively, the sorting theorem simplifies to

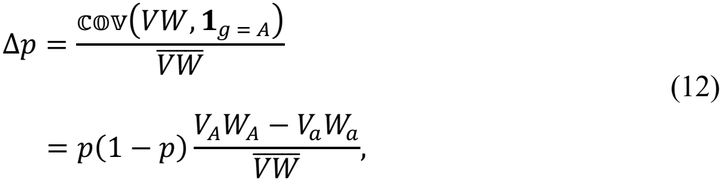

where 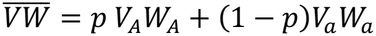. Equation (12) recovers the haploid expansion model of Phillips and Perkins (2019), here obtained as a special case of the sorting theorem.

#### Diploid model of gene evolution

Consider an infinitely large population of diploid parents with genotypes *AA, Aa*, and *aa* and sorting fitness *V*_*AA*_*W*_*AA*_, *V*_*Aa*_*W*_*Aa*_, and *V*_*aa*_*W*_*aa*_, respectively. Under Hardy-Weinberg equilibrium, the genotype frequencies are *p*^2^, 2*p*(1 − *p*), and (1 − *p*)^2^, respectively, where *p* is the frequency of allele *A* in the population. We define *i*’s phenotype as

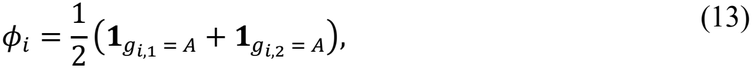

which corresponds to the frequency of allele *A* in *i*’s genotype, with *g*_*i,j*_ representing the identity of the *j*th allele copy. Thus, the phenotype of the *i*th parent, *ϕ*_*i*_, can take values 1, 1/2, and 0 for genotypes *AA, Aa*, and *aa*, respectively. As in the haploid model, with no mutation, the evolutionary change is given by

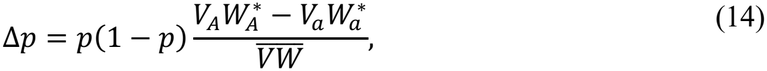

where 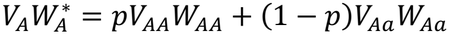 and 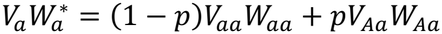 are the *marginal sorting fitness* of alleles *A* and *a*, respectively, such that 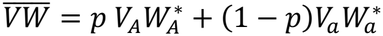. This recovers the diploid expansion model of Phillips and Perkins (2019).

#### Haploid models of gene surfing

To derive the models of gene surfing, we adapt the haploid model of gene evolution by Phillips and Perkins (2019) with three modifications. We assume that the realized sorting fitness of a parent is a random variable drawn from a probability distribution, no parent has a reproductive or dispersal advantage, and the population size of offspring in patch *x*′ is constant and is equal to *N*. These assumptions allow several mathematical simplifications. We can set 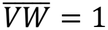, which in turn implies that 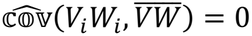. Simplifying this constraint yields

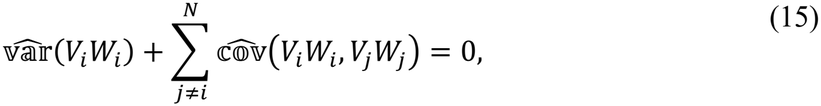

where 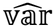 and 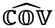 are the variance and covariance of random variables. We can assume that *V*_*i*_*W*_*i*_ are exchangeable random variables. Consequently, we can set 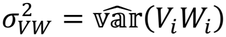 and 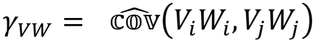, which further simplifies equation (15) to 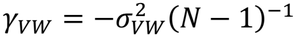. Using the sorting theorem (with 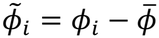) and the mathematical results described above, we can show that the variance of the change in allele frequency is

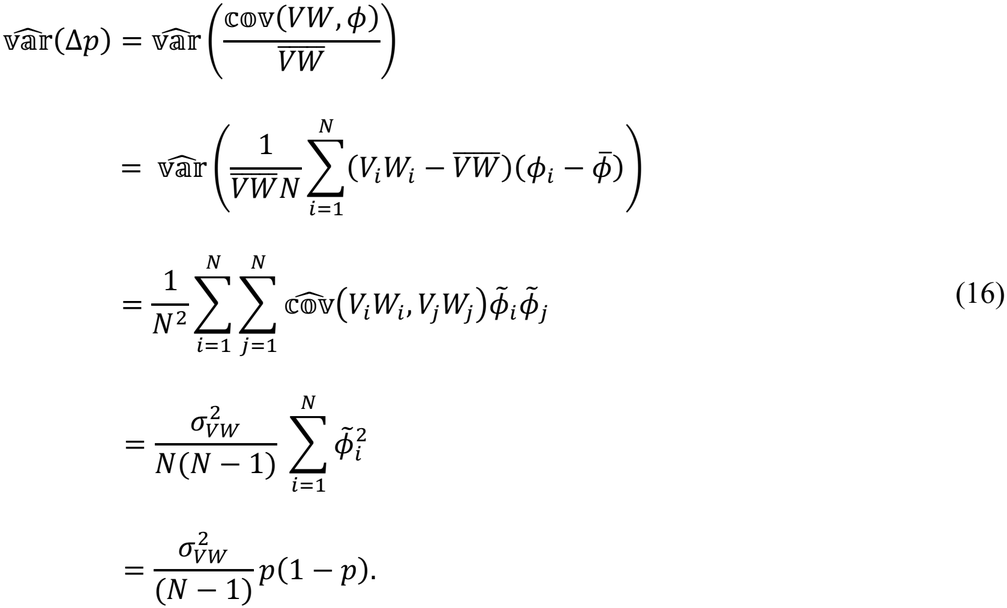

To calculate 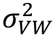, we consider two scenarios. First, we assume that parents in patch *x* produce an infinitely large number of offspring, of which *N* colonize patch *x′*. Because each offspring in patch *x′* has a probability *N*^−1^ that its parent is *i*, the total number of offspring of *i* in patch *x′, V*_*i*_*W*_*i*_, is a Binomial(*N, N*^−1^) random variable with variance 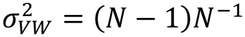. Consequently, the variance in the change in allele frequency is

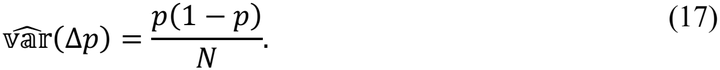

Equation (17) is the variance result for gene surfing that was derived in Peischl and Gilbert (2020). Second, we can consider an alternative setup in which *k* parents are randomly selected from patch *x*, such that each selected parent leaves *N*/*k* offspring in patch *x′*. Under these assumptions, we can show that 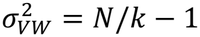, which under the infinitely large population limit (i.e., *N* ⟶ ∞) gives the variance result for haploid individuals from Slatkin and Excoffier (2012),

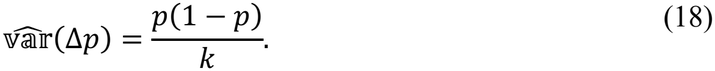

#### Quantitative genetics model of phenotypic evolution

Consider an infinitely large population of parents with a continuously varying phenotype *ϕ*_*i*_. We assume that an individual’s phenotype is controlled by both environmental factors and a large number of genes, each contributing a small effect. Assuming that the joint phenotypic distribution of parent and offspring follows a multivariate normal distribution, we can express the mean offspring phenotype as a linear function of the parents’ phenotype,

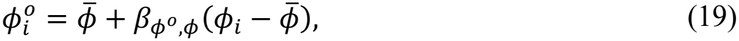

where 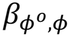 is the slope of the regression line between mean offspring and parents’ phenotypes. This slope is commonly referred to as the heritability (*h*^2^) and is the ratio of additive genetic variance (*G*) to phenotypic variance (var(*ϕ*) = *P*). Substituting this inheritance relationship into the sorting theorem gives 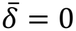 and

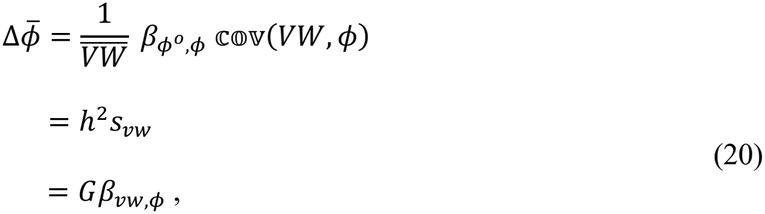

where *s*_*vw*_ is the *sorting differential*, and *β*_*vw,ϕ*_ is the *sorting gradient* (slope of the regression line between 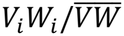 and *ϕ*_*i*_). Goel et al. (2025) derived this model to measure the strength of phenotypic evolution in a moving frame of reference.

#### Multivariate quantitative genetics model of phenotypic evolution

To derive the multivariate extension of equation (20), we make a few additional assumptions. We assume that the sorting fitness and mean offspring phenotypes are linear functions of *n* continuously varying parental phenotypes, denoted by a vector 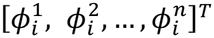. As before, these phenotypes are determined by environmental factors and the action of multiple genes. However, a gene can control multiple phenotypes due to physical linkage and pleiotropy (Lande and Arnold 1983; Walsh and Lynch 2018). Consequently, a phenotype can evolve through its direct effect on the individual’s sorting fitness as well as through indirect effects via genetic correlations with other phenotypes. Using path analysis (Rice 2004), we can show that

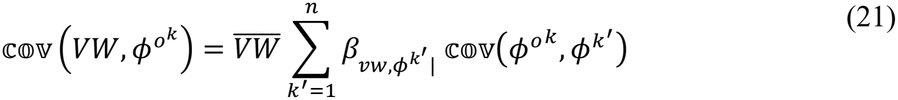

where 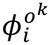 is the mean value of the *k*th phenotype of offspring of the *i*th parent, and 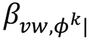 is the partial regression of 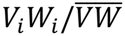 on 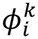. Equation (21) captures both the linear inheritance relationship and the causal link between parents’ phenotype and sorting fitness. By substituting equation (21) into the sorting theorem corresponding to each trait, we can show that the evolutionary change is given by

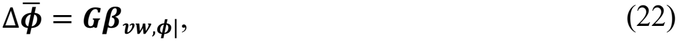

where 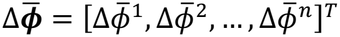, ***G*** is the additive genetic covariance matrix (or the *G*-matrix for short), and 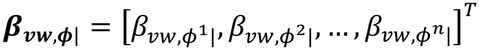. As in the single-trait quantitative genetics model of Goel et al. (2025), the partial sorting gradients 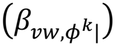 quantify the magnitude of phenotypic change when multiple traits jointly govern evolution.

These derivations highlight that the sorting theorem is indeed a general descriptor of evolution in a moving frame of reference. We show that despite the differences in directional and non-directional expansion models of evolution (Eqs. 12, 14, 17, 18, 20, and 22), the sorting theorem reveals them as specific instances of a common mathematical framework.

### Implications

Since Price’s theorem is a general descriptor of both classical and expansion models of evolution, using Price’s theorem is a better way to evaluate whether spatial sorting and gene surfing are spatial analogs of selection and genetic drift, respectively, than the current approach of comparing special-case models one by one (for example, see Peischl and Gilbert 2020; Phillips and Perkins 2019; Slatkin and Excoffier 2012). This is because, by using Price’s theorem as the starting point, we can be reasonably confident that our inferences apply to a broad class of analogous biological systems, and we are not misled by model-specific assumptions.

We find that gene surfing is indeed a spatial analog of genetic drift. Price’s theorem yields the standard variance result for genetic drift and gene surfing when fitness and sorting fitness are treated as random variables. However, the analogy breaks down when we compare directional models of evolution. Spatial sorting, as defined by Shine et al. (2011), refers specifically to the evolution of dispersal in the absence of differential lifetime reproductive success. In contrast, the sorting theorem shows that directional evolution in a moving frame of reference is driven by the number of offspring a parent leaves in a newly colonized patch, which depends on both non-random dispersal and reproduction. This finding results in a series of conceptual challenges.

To shed light on these challenges, we must first understand why spatial sorting rose to prominence. A primary requirement for selection is differential lifetime reproductive success (Darwin 1859). However, recent empirical and theoretical work showed that at the invasion front, dispersal-related traits could evolve even without differential fitness, thereby identifying a novel evolutionary mechanism that cannot be explained by classical evolutionary theory. This evolutionary process was formalized (often verbally with simulations) in a series of conceptual papers (Perkins et al. 2013; Phillips et al. 2010; Shine et al. 2011; Travis and Dytham 2002). However, these papers made two important claims that are rarely discussed in the literature but have important conceptual implications.

First, the papers on expansion models propose a novel approach for measuring evolutionary change by tracking changes in mean phenotype in a moving frame of reference. To understand its importance, consider Price’s theorem. We show that in a static and moving frame of reference, the *w*_*i*_ term in Price’s theorem corresponds to fitness (*W*_*i*_) and sorting fitness (*V*_*i*_*W*_*i*_), respectively. Therefore, in a moving frame of reference, evolution of dispersal in the absence of differential lifetime reproductive success is simply an outcome of how we choose to measure evolutionary change. We arrive at this inference from Price’s theorem, which is a general mathematical descriptor of evolution, rather than from a model-by-model comparison.

Second, papers on expansion models often use terminology that overlaps with classical models of evolution. For example, Phillips et al. (2010) define ‘spatial selection’ as the evolution of dispersal at the population vanguard due to spatial assortment of dispersal phenotypes (spatial sorting) and improved fitness at low-density edge populations (natural selection). Perkins et al. (2013) define ‘enhanced spatial selection’ as a scenario in which evolution of dispersal due to spatial selection is enhanced by evolution of other life history traits. These definitions create a conceptual problem—should we choose terminology based on what drives evolution or how evolution is measured? To illustrate this problem, consider two expansion models of evolution: the haploid model by Phillips and Perkins (2019) with a constant value of *V*_*i*_ for all parents,

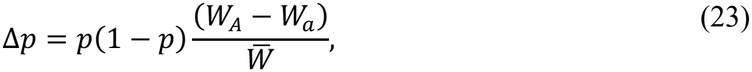

and the haploid model of gene surfing (Eq. 17). By current convention, equation (23) is considered a model of natural selection because evolution is driven by differential lifetime reproductive success, and the equation of evolution matches the haploid model of gene selection by Haldane (1924). However, the variance result of gene surfing (Eq. 17) also matches exactly with the variance result for genetic drift (Wright 1931), yet they are treated as distinct evolutionary processes because of how allele frequency change is measured (Peischl and Gilbert 2020). To resolve these terminological inconsistencies, we see three possibilities.

- We expand the definition of selection to encompass any evolutionary process that produces directional phenotypic change, irrespective of how that change is measured. Under this proposal, evolutionary change in a moving frame of reference, driven by differential dispersal capacity, differential lifetime reproductive success, or both, would be treated as natural selection. Consequently, spatial sorting (as defined in Shine et al. 2011), spatial selection (Phillips et al. 2010), and enhanced spatial selection (Perkins et al. 2013) would be reclassified as natural selection. Furthermore, to maintain consistency, we must also expand the definition of genetic drift to encompass non-directional evolution in both static and moving frames of reference. As a result, gene surfing (as defined in Klopfstein et al. 2006) would be reclassified as genetic drift.
- We define a new term, spatial assortment, for describing directional evolution in a moving frame of reference. Under this proposal, evolutionary change in a moving frame of reference, driven by differential dispersal capacity, differential lifetime reproductive success, or both, would be treated as spatial assortment. Consequently, spatial sorting, spatial selection, and enhanced spatial selection would all be considered as special cases of spatial assortment. Furthermore, spatial assortment, not spatial sorting, would be considered as the spatial analog of selection. Both gene surfing and genetic drift would retain their current definitions.
- Lastly, we expand the definition of spatial sorting to mean directional evolutionary change in a moving frame of reference. Under this proposal, evolutionary change in a moving frame of reference driven by differential dispersal capacity, differential lifetime reproductive success, or both, would be treated as spatial sorting. Both gene surfing and genetic drift would retain their current definitions. We advocate for the third proposal for two reasons. First, the third proposal recognizes that measuring evolution in a moving frame of reference offers a new vantage point for describing natural variation. Second, with the expanded definition of spatial sorting, we can correctly label it as the spatial analog of natural selection.

The sorting theorem can also serve other purposes, such as organizing our evolutionary reasoning, identifying connections across models of evolution, critiquing theoretical arguments, and, most importantly, building a unified theory of expansion models. Unlike classical evolutionary theory, the mathematical theory of spatial sorting is still in its infancy. The sorting theorem can help researchers jump-start theory-building by stimulating new ideas and mathematical models, which can, in turn, spark new empirical investigations.

## Acknowledgments

I acknowledge the support from the Continuing and Stengl-Wyer fellowships by The University of Texas at Austin. I would like to thank Stephen Stearns, Arun Chavan, Timothy Keitt, Renee Duckworth, Athmanathan Senthilnathan, Ben Phillips, Sean Rice, Damla Cinoglu, Mattheau Comerford, Scott Egan, and Thomas Juenger for their feedback.

