## Supplementary Information for "Evolution in a moving frame of reference"

We present our mathematical derivations of directional and non-directional models of evolution that quantify phenotypic changes in a moving frame of reference. All models are derived starting from the sorting theorem by supplying simplifying assumptions. These derivations closely follow how Price's theorem has been used to derive directional and non-directional models of classical evolution (Rice 2004).

*Haploid model of gene evolution by Phillips and Perkins (2019)*

Consider an infinitely large population of haploid parents with genotypes $A$ and $a$ with frequency $p$ and $1-p$, respectively. Thus, the numbers of haploid individuals with genotypes $A$ and $a$ are given by $Np$ and $N(1-p)$, respectively. We define the phenotype of the $i$th parent as

$$\phi_i = \mathbf{1}_{g_i = A}, \tag{A1}$$

which equals one if the $i$th parent has genotype $g_i = A$ and zero if $g_i = a$. By construction,

$$\begin{aligned} \bar{\phi} &= \frac{\sum_{k=1}^{Np} 1 + \sum_{k=1}^{N(1-p)} 0}{N} \\ &= \frac{Np}{N} \\ &= p, \end{aligned} \tag{A2}$$

such that the evolutionary change $\Delta p$ is the difference between the frequency of genotype $A$ among offspring in patch $x'$ and $\bar{\phi}$. Assuming perfect transmission (no mutation, so $\bar{\delta} = 0$ and $\phi_i^o = \phi_i$) and denoting the sorting fitness of parents with genotypes $A$ and $a$ by $V_A W_A$ and $V_a W_a$, respectively, the sorting theorem simplifies to

$$\begin{aligned} \Delta p &= \frac{\mathbb{cov}(VW, \phi)}{\overline{VW}} \\ &= \frac{\overline{VW\phi} - \overline{VW}\bar{\phi}}{\overline{VW}} \\ &= \frac{\overline{VW\phi} - \overline{VW}p}{\overline{VW}}. \end{aligned} \tag{A3}$$

Here, the mean population sorting fitness is given by

$$\begin{aligned} \overline{VW} &= \frac{\sum_{k=1}^{Np} V_A W_A + \sum_{k=1}^{N(1-p)} V_a W_a}{N} \\ &= p\, V_A W_A + (1-p) V_a W_a \end{aligned} \tag{A4}$$

and

$$\overline{VW\phi} = \frac{\sum_{k=1}^{Np} V_A W_A \,.\, 1 + \sum_{k=1}^{N(1-p)} V_a W_a . \, 0}{N} \\ = pV_A W_A. \tag{A5}$$

Substituting A4 and A5 in A3, we get

$$\Delta p = \frac{pV_A W_A - (p^2\, V_A W_A + p(1-p)V_a W_a)}{\overline{VW}} \\ = p(1-p)\frac{V_A W_A - V_a W_a}{\overline{VW}}. \tag{A6}$$

*Diploid model of gene evolution by Phillips and Perkins (2019)*

Consider an infinitely large population of diploid parents with genotypes $AA$, $Aa$, and $aa$ with sorting fitness $V_{AA}W_{AA}$, $V_{Aa}W_{Aa}$, and $V_{aa}W_{aa}$, respectively. Under the Hardy-Weinberg equilibrium, the frequencies of these genotypes are $p^2$, $2p(1-p)$, and $(1-p)^2$, respectively, where $p$ is the frequency of allele $A$ in the population. We define $i$'s phenotype as

$$\phi_i = \frac{1}{2}\left(\mathbf{1}_{g_{i,1} = A} + \mathbf{1}_{g_{i,2} = A}\right), \tag{A7}$$

which corresponds to the frequency of the allele $A$ in $i$'s genotype, with $g_{i,j}$ representing the identity of the $j$th copy of the allele. Since the individuals are diploid, $j$ takes values one and two. Thus, the phenotype of the $i$th parent, $\phi_i$, can take values one, half, and zero for genotypes $AA$, $Aa$, and $aa$, respectively. The mean phenotype of parents in the population is given by

$$\bar{\phi} = \frac{\sum_{k=1}^{Np^2} 1 + \sum_{k=1}^{2Np(1-p)} 1/2 + \sum_{k=1}^{N(1-p)^2} 0}{N} \\ = \frac{N\left(p^2 + p(1-p)\right)}{N} \\ = p. \tag{A8}$$

As in the haploid model, with no mutation, the evolutionary change is given by

$$\Delta p = \frac{\mathbb{cov}(VW, \phi)}{\overline{VW}} \\ = \frac{\overline{VW\phi} - \overline{VW}\bar{\phi}}{\overline{VW}} \\ = \frac{\overline{VW\phi} - \overline{VW}p}{\overline{VW}}. \tag{A9}$$

For a diploid population, the mean population sorting fitness is given by

$$\overline{VW} = \frac{\sum_{k=1}^{Np^2} V_{AA}W_{AA} + \sum_{k=1}^{2Np(1-p)} V_{Aa}W_{Aa} + \sum_{k=1}^{N(1-p)^2} V_{aa}W_{aa}}{N} \qquad \text{(A10)}$$
$$= p^2\, V_{AA}W_{AA} + 2p(1-p)V_{Aa}W_{Aa} + (1-p)^2\, V_{aa}W_{aa}$$

and

$$\overline{VW\phi} = \frac{\sum_{k=1}^{Np^2} V_{AA}W_{AA}.1 + \sum_{k=1}^{2Np(1-p)} V_{Aa}W_{Aa}\,.1/2 + \sum_{k=1}^{N(1-p)^2} V_{aa}W_{aa}.0}{N} \qquad \text{(A11)}$$
$$= p^2V_{AA}W_{AA} + p(1-p)V_{Aa}W_{Aa}.$$

Defining the marginal sorting fitness of alleles $A$ and $a$ as

$$V_AW_A^* = pV_{AA}W_{AA} + (1-p)V_{Aa}W_{Aa} \qquad \text{(A12)}$$

and

$$V_aW_a^* = (1-p)V_{aa}W_{aa} + pV_{Aa}W_{Aa}, \qquad \text{(A13)}$$

respectively, we can express $\overline{VW}$ and $\overline{VW\phi}$ as

$$\overline{VW} = pV_AW_A^* + (1-p)V_aW_a^* \qquad \text{(A14)}$$

and

$$\overline{VW\phi} = pV_AW_A^*. \qquad \text{(A15)}$$

Substituting A14 and A15 in A9, we get

$$\Delta p = \frac{pV_AW_A^* - (p^2\, V_AW_A^* + p(1-p)V_aW_a^*)}{\overline{VW}} \qquad \text{(A16)}$$
$$= p(1-p)\frac{V_AW_A^* - V_aW_a^*}{\overline{VW}}.$$

### *Haploid models of gene surfing by Peischl and Gilbert (2020) and Slatkin and Excoffier (2012)*

To derive the models of gene surfing by Peischl and Gilbert (2020) and Slatkin and Excoffier (2012), we adapt the haploid model of gene evolution by Phillips and Perkins (2019) with three modifications. We assume that the realized sorting fitness of a parent is a random variable drawn from a probability distribution, no parent has a reproductive or dispersal advantage, and the population size of offspring in patch $x'$ is finite and is equal to $N$. These assumptions allow several mathematical simplifications. First, we can set $\overline{VW} = 1$, which in turn implies that $\widehat{\mathrm{cov}}(V_iW_i, \overline{VW}) = 0$. Simplifying this constraint yields

$$\widehat{\mathbb{cov}}\left(V_iW_i, N^{-1}\sum_{j=1}^{N} V_jW_j\right) = 0$$
$$\sum_{j=1}^{N} \widehat{\mathbb{cov}}\big(V_iW_i, V_jW_j\big) = 0 \tag{A17}$$
$$\widehat{\mathbb{var}}(V_iW_i) + \sum_{j\neq i}^{N} \widehat{\mathbb{cov}}\big(V_iW_i, V_jW_j\big) = 0,$$

where $\widehat{\mathbb{var}}$ and $\widehat{\mathbb{cov}}$ are the variance and covariance of random variables. Second, we can assume that the sorting fitnesses of parents are exchangeable random variables. Intuitively, the exchangeability implies that the joint distribution of $V_1W_1$, $V_2W_2$, … , $V_NW_N$ is unchanged when the labels of any two parents $i$ and $j$ are swapped:

$$\begin{aligned} &P\big[V_1W_1 = k_1, \dots, V_iW_i = k_i, \dots, V_jW_j = k_j, \dots, V_NW_N = k_N | \overline{VW} = 1\big] \\ &\quad = P\big[V_1W_1 = k_1, \dots, V_iW_i = k_j, \dots, V_jW_j = k_i, \dots, V_NW_N = k_N | \overline{VW} \\ &\quad = 1\big]. \end{aligned} \tag{A18}$$

This allows us to set $\sigma_{VW}^2 = \widehat{\mathbb{var}}(V_iW_i)$ and $\gamma_{VW} = \widehat{\mathbb{cov}}\big(V_iW_i, V_jW_j\big)$, which further simplifies the equation (A17) to $\gamma_{VW} = -\sigma_{VW}^2(N-1)^{-1}$.

Using the sorting theorem (with $\tilde{\phi}_i = \phi_i - \bar{\phi}$, such that $\sum_{i=1}^{N} \tilde{\phi}_i = 0$), the mathematical results described above, and noting that $\phi_i^2 = \phi_i$, we can show that the variance of the change in allele frequency is

$$\begin{aligned} \widehat{\mathbb{var}}(\Delta p) &= \widehat{\mathbb{var}}\left(\frac{\mathbb{cov}(VW, \phi)}{\overline{VW}}\right) \\ &= \widehat{\mathbb{var}}\left(\frac{1}{\overline{VW}N}\sum_{i=1}^{N}(V_iW_i - \overline{VW})(\phi_i - \bar{\phi})\right) \\ &= \frac{1}{N^2}\widehat{\mathbb{var}}\left(\sum_{i=1}^{N} V_iW_i\tilde{\phi}_i\right) \\ &= \frac{1}{N^2}\sum_{i=1}^{N}\sum_{j=1}^{N}\widehat{\mathbb{cov}}\big(V_iW_i, V_jW_j\big)\tilde{\phi}_i\tilde{\phi}_j \\ &= \frac{1}{N^2}\sum_{i=1}^{N}\widehat{\mathbb{var}}(V_iW_i)\tilde{\phi}_i^2 + \frac{1}{N^2}\sum_{i=1}^{N}\sum_{j\neq i}^{N}\widehat{\mathbb{cov}}\big(V_iW_i, V_jW_j\big)\tilde{\phi}_i\tilde{\phi}_j \end{aligned}$$

$$= \frac{1}{N^2}\left(\sigma_{VW}^2 \sum_{i=1}^{N} \tilde{\phi}_i^2 + \gamma_{VW} \sum_{i=1}^{N} \sum_{j \neq i}^{N} \tilde{\phi}_i \tilde{\phi}_j\right) \tag{A19}$$

$$= \frac{1}{N^2}\left(\sigma_{VW}^2 \sum_{i=1}^{N} \tilde{\phi}_i^2 + \gamma_{VW}\left(\left(\sum_{i=1}^{N} \tilde{\phi}_i\right)^2 - \sum_{i=1}^{N} \tilde{\phi}_i^2\right)\right)$$

$$= \frac{1}{N^2}(\sigma_{VW}^2 - \gamma_{VW}) \sum_{i=1}^{N} \tilde{\phi}_i^2$$

$$= \frac{1}{N}\frac{\sigma_{VW}^2}{(N-1)} \sum_{i=1}^{N} (\phi_i^2 - 2\phi_i\bar{\phi} + \bar{\phi}^2)$$

$$= \frac{1}{N}\frac{\sigma_{VW}^2}{(N-1)} (N\bar{\phi} - 2N\bar{\phi}\bar{\phi} + N\bar{\phi}^2)$$

$$= \frac{\sigma_{VW}^2}{(N-1)} p(1-p)$$

To calculate $\sigma_{VW}^2$, we consider two scenarios. First, we assume that parents in patch $x$ produce an infinitely large number of offspring, of which $N$ colonize patch $x'$. Because each offspring in patch $x'$ has a probability $N^{-1}$ that its parent is $i$, the total number of offspring of $i$ in patch $x'$, $V_i W_i$, is a Binomial random variable with probability of success $N^{-1}$ and number of trials $N$. Consequently, $\sigma_{VW}^2 = (N-1)N^{-1}$, and the variance in the change in allele frequency is

$$\widehat{\mathbb{var}}(\Delta p) = \frac{p(1-p)}{N}. \tag{A20}$$

This variance result matches the result of gene surfing by Peischl and Gilbert (2020). Second, we consider that patch $x'$ is founded by $k$ randomly selected parents, and each founding parent leaves $N/k$ offspring. Consequently, $\widehat{\mathbb{E}}((V_i W_i)^2) = (N/k)^2\,(k/N) + 0^2(1 - k/N) = N/k$ ($\widehat{\mathbb{E}}$ is the expectation value of the random variable) and $\sigma_{VW}^2 = N/k - 1$. Thus, we can show the variance in the change in allele frequency is

$$\widehat{\mathbb{var}}(\Delta p) = \frac{N-k}{N-1}\frac{p(1-p)}{k}. \tag{A21}$$

In the limit of infinitely large population (i.e., $N \longrightarrow \infty$), we recover the variance result from Slatkin and Excoffier (2012)

$$\widehat{\mathbb{var}}(\Delta p) = \frac{p(1-p)}{k}. \tag{A22}$$

This variance result is missing a factor of ½ (see eq. 2 in Appendix A of Slatkin and Excoffier 2012) as we consider haploid individuals.

*Quantitative genetics model of phenotypic evolution by Goel et al. (2025)*

Consider an infinitely large population of parents with a continuously varying phenotype $\phi_i$. We assume that an individual's phenotype is controlled by both environmental factors and a large number of genes, each contributing a small effect. Assuming that the joint phenotypic distribution of parent and offspring follows a multivariate normal distribution, we can express the mean offspring phenotype as a linear function of the parents' phenotype,

$$\phi_i^o = \bar{\phi} + \beta_{\phi^o,\phi}(\phi_i - \bar{\phi}), \tag{A23}$$

where $\beta_{\phi^o,\phi}$ is the slope of the regression line between mean offspring and parents' phenotypes (Falconer 1960). This slope is commonly referred to as the heritability ($h^2$) and is the ratio of additive genetic variance (*G*) to phenotypic variance ($\mathbb{var}(\phi) = P$). Substituting this inheritance relationship in the sorting theorem gives

$$\begin{aligned}\bar{\delta} &= \frac{1}{N}\sum_{i=1}^{N}\left(\beta_{\phi^o,\phi} - 1\right)(\phi_i - \bar{\phi}) \\ &= \left(\beta_{\phi^o,\phi} - 1\right)\left(\left(\sum_{i=1}^{N}\frac{\phi_i}{N}\right) - \bar{\phi}\right) \\ &= 0\end{aligned} \tag{A24}$$

and

$$\begin{aligned}\Delta\bar{\phi} &= \frac{1}{\overline{\overline{VW}}}\ \mathbb{cov}\left(VW, \beta_{\phi^o,\phi}(\phi - \bar{\phi})\right) \\ &= \frac{1}{\overline{\overline{VW}}}\ \beta_{\phi^o,\phi}\ \mathbb{cov}(VW, \phi) \\ &= \frac{G}{P}\ \beta_{vw,\phi}\mathbb{var}(\phi) \\ &= G\beta_{vw,\phi}\end{aligned} \tag{A25}$$

Here, $\beta_{vw,\phi}$ (sorting gradient) is the slope of the regression line between $V_iW_i/\overline{VW}$ and $\phi_i$.

*Multivariate quantitative genetics model of phenotypic evolution*

To derive the multivariate quantitative genetics model, we assume that the sorting fitness and mean offspring phenotypes are linear functions of $n$ continuously varying parental phenotypes, denoted by a vector $[\phi_i^1,\ \phi_i^2, \ldots, \phi_i^n]^T$. We assume that the distribution of parent and offspring traits follows a multivariate normal distribution. For simplicity, we first derive the result for $n = 2$ and then generalize to an arbitrary number of traits. Using path analysis, we can show

$$\frac{1}{\overline{VW}}\mathbb{cov}\left(VW, \phi^{o^1}\right) = \begin{array}{l} \beta_{vw,\phi^1|}\mathbb{cov}(\phi^1,\phi^1)\beta_{\phi^{o^1},\phi^1|} + \beta_{vw,\phi^1|}\mathbb{cov}(\phi^1,\phi^2)\beta_{\phi^{o^1},\phi^2|} \\ +\beta_{vw,\phi^2|}\mathbb{cov}(\phi^2,\phi^2)\beta_{\phi^{o^1},\phi^2|} + \beta_{vw,\phi^2|}\mathbb{cov}(\phi^1,\phi^2)\beta_{\phi^{o^1},\phi^1|} \end{array} \tag{A26}$$

and

$$\begin{aligned} \mathbb{cov}\left(\phi^{o^1},\phi^1\right) &= \mathbb{cov}(\phi^1,\phi^1)\beta_{\phi^{o^1},\phi^1|} + \mathbb{cov}(\phi^1,\phi^2)\beta_{\phi^{o^1},\phi^2|} \\ \mathbb{cov}\left(\phi^{o^1},\phi^2\right) &= \mathbb{cov}(\phi^2,\phi^2)\beta_{\phi^{o^1},\phi^2|} + \mathbb{cov}(\phi^1,\phi^2)\beta_{\phi^{o^1},\phi^1|}, \end{aligned} \tag{A27}$$

where the '|' sign in the subscript of $\beta_{y,x|}$ is shorthand for the partial regression of $y$ on $x$. Combining equations (A23) and (A24), we get

$$\frac{1}{\overline{VW}}\mathbb{cov}\left(VW, \phi^{o^1}\right) = \beta_{vw,\phi^1|}\ \mathbb{cov}\left(\phi^{o^1},\phi^1\right) + \beta_{vw,\phi^2|}\ \mathbb{cov}\left(\phi^{o^1},\phi^2\right). \tag{A28}$$

Similarly, we can write an analogous equation for $\mathbb{cov}\left(VW, \phi^{o^2}\right)$. Using the sorting theorem, we can show $\bar{\delta}^1 = \bar{\delta}^2 = 0$ and

$$\begin{aligned} \Delta\bar{\phi}^1 &= \beta_{vw,\phi^1|}\ \mathbb{cov}\left(\phi^{o^1},\phi^1\right) + \beta_{vw,\phi^2|}\ \mathbb{cov}\left(\phi^{o^1},\phi^2\right) \\ \Delta\bar{\phi}^2 &= \beta_{vw,\phi^1|}\ \mathbb{cov}\left(\phi^{o^2},\phi^1\right) + \beta_{vw,\phi^2|}\ \mathbb{cov}\left(\phi^{o^2},\phi^2\right). \end{aligned} \tag{A29}$$

By extension, for an arbitrary number of traits

$$\Delta\bar{\phi}^k = \sum_{k'=1}^{n}\beta_{vw,\phi^{k'}|}\ \mathbb{cov}\left(\phi^{o^k},\phi^{k'}\right) \tag{A30}$$

and $\bar{\delta}^k = 0$. In matrix form, equation (A27) can be re-expressed as

$$\begin{bmatrix}\Delta\bar{\phi}^1 \\ \Delta\bar{\phi}^2 \\ \vdots \\ \Delta\bar{\phi}^n\end{bmatrix} = \begin{bmatrix} \mathbb{cov}\left(\phi^{o^1},\phi^1\right) & \mathbb{cov}\left(\phi^{o^1},\phi^2\right) & \ldots & \mathbb{cov}\left(\phi^{o^1},\phi^n\right) \\ \mathbb{cov}\left(\phi^{o^2},\phi^1\right) & \mathbb{cov}\left(\phi^{o^2},\phi^2\right) & \ldots & \mathbb{cov}\left(\phi^{o^2},\phi^n\right) \\ \vdots & \vdots & & \vdots \\ \mathbb{cov}\left(\phi^{o^n},\phi^1\right) & \mathbb{cov}\left(\phi^{o^n},\phi^2\right) & \ldots & \mathbb{cov}\left(\phi^{o^n},\phi^n\right) \end{bmatrix} \begin{bmatrix}\beta_{vw,\phi^1|} \\ \beta_{vw,\phi^2|} \\ \vdots \\ \beta_{vw,\phi^n|}\end{bmatrix}, \tag{A31}$$

which can be succinctly represented as

$$\Delta\bar{\boldsymbol{\phi}} = \boldsymbol{G}\boldsymbol{\beta}_{\boldsymbol{vw},\boldsymbol{\phi}|}. \tag{A32}$$

Here, $\boldsymbol{G}$ is the additive genetic covariance matrix (Lande and Arnold 1983).
